# The *Pseudogymnoascus destructans* Proteome Under Copper Stress Conditions

**DOI:** 10.64898/2026.03.13.711597

**Authors:** Alyssa D. Friudenberg, Saika Anne, Yuan Lu, Susan T. Weintraub, Ryan L. Peterson

## Abstract

The invasive fungal pathogen *Pseudogymnoascus destructans* is responsible for the collapse of several North American bat species through an infectious fungal skin disease known as White-Nose Syndrome (WNS). Recent transcriptomic studies have suggested that trace copper ion acquisition is essential for *P. destructans* propagation on its animal hosts. However, little is known about the mechanistic details of *P. destructans* adaptation occurring at the protein level. In this study, we report the global proteomic adaptation of *P. destructans* under chronic Cu-stress growth conditions employing chemically defined media. We identify 4340 *P. destructans* proteins, or approximately 47.8% of the predicted proteome, spanning a dynamic intensity range of six orders of magnitude. Chronic Cu-withholding stress leads to substantial alterations in the proteome, with 1398 differentially abundant proteins (DAPs) exhibiting statistically significant (p < 0.05) changes in protein levels compared to control growth conditions. We find that Cu-withholding stress induces increased levels of proteins associated with high-affinity Cu-acquisition, changes in intracellular superoxide dismutase (SOD) levels, and alterations in mitochondrial proteins related to aerobic respiration. In contrast, chronic Cu-overload stress leads to 390 DAPs (p < 0.05), which are more widely distributed across the proteome, with several DAPs associated with genomic stability and basic metabolism. Additionally, in this report, we present assessment of antisera products against intracellular and cell-surface protein targets of *P. destructans* that are effective for indicating Cu-withholding stress by western blotting.

## 1. Introduction

The opportunistic fungal pathogen *Pseudogymnoascus destructans* causes the infectious skin disease White-Nose Syndrome (WNS) and has led to the decimation of several North American cave-dwelling bat populations^1–3^. *P. destructans* has a unique evolutionary trajectory as an animal-infecting fungal pathogen. It is hypothesized that *P. destructans* originated as a soil-dwelling plant-infecting pathogen that adapted to thrive on bat animal hosts^4^. The ability of *P. destructans* to evolve mechanisms that enable it to thrive across diverse ecological niches, including soil, cave hibernacula, plants, and animal hosts, suggests that studying this fungal pathogen may provide insights into host-microbe interactions that enhance the pathogenicity of environmental microbes^5–7^. Similarities between WNS disease pathology found in post-hibernation emergent euthermic bats and immune reconstitution inflammatory syndrome (IRIS) found in humans,^8^ provide support that further investigation of *P. destructans* and WNS dynamics will yield insights into a better understanding of how fungal burden and fungal secreted factors contribute to this complex immune response in animal hosts.

To efficiently colonize and propagate on a vulnerable animal host, fungal pathogens must evade the host immune response^9^ and harvest essential micronutrients, including trace metals, from their host to maintain basic metabolic processes^10^. Through a process known as nutritional immunity,^11^ animal hosts utilize high-affinity metal-binding proteins and sequestration strategies to restrict microbe access to trace metal resources^12^. Alternatively, toxic levels of copper can be delivered into the phagolysosome to boost the respiratory burst killing efficiency^13, 14^. The ability to navigate extremes in copper bioavailability through high-affinity copper import and copper efflux is vital for fungal virulence^15^.

Copper, as a trace micronutrient in fungi, participates as a required cofactor for several metalloenzymes, including Cu-containing superoxide dismutases (SODs), cytochrome *c* oxidase (CcO), and multicopper oxidases (MCOs)^16, 17^. Together, these cuproenzymes assist in maintaining intracellular redox homeostasis and participate in forming a protective barrier from external insults enzymatically through SOD activity or physically via melanin^18–20^. At active WNS fungal-infection sites, elevated bat-host S100 protein transcripts^21^ and *P. destructans* high-affinity Cu-transporter (CTR) gene transcripts^22^ can be detected. These observations suggest that copper bioavailability is low at the host-pathogen interface, and the bat-host is restricting metal nutrients to circumvent fungal growth. Indeed, *in vitro* transcriptomic studies that systemically restricted Cu bioavailability found a similar *P. destructans* transcriptional response to that observed on the bat host, including the regulation of several putative copper-utilizing virulence factors associated with WNS fungal pathology^23^. Furthermore, recent studies with *P. destructans* conidia have demonstrated that fungal melanin can also promote intracellular invasion and gemination in bat keratinocytes^24^. Together, these transcriptional studies suggest that copper dynamics at the host-pathogen interface are relevant to WNS disease propagation, and that there is a battle over trace copper resources between the bat host and the *P. destructans* fungal pathogen.

While transcriptional methods offer powerful insights into gene expression profiles under environmental stress, protein levels are not always correlated with transcript levels. In a previous *in vitro* study investigating the *P. destructans* Cu-stress transcriptional response by our lab,^23^ we identified a collection of genes that display reciprocal regulation in response to Cu-withholding and Cu-overload stress, which we hypothesize participate in two putative high-affinity Cu-import pathways. The first pathway involves the canonical family of high-affinity Cu-transporters and multiple isoforms related to the recently identified secreted Cu-scavenging protein BIM1/Cbi1 found in *Cryptococcus neoformans*^25, 26^. The second putative Cu-scavenging pathway, identified as the Cu-responsive gene Cluster (CRC), may involve a small-molecule metallophore^23^. The CRC encodes for genes homologous to the biosynthetic pathway found in *Staphylococcus aureus*^27^ and *Pseudomonas aeruginosa*^28^, which encode for metal-binding opine-metallophores. Two important questions arise regarding the two possible identified Cu-trafficking pathways: Does *P. destructans* express multiple BIM1/Cbi1 isoforms in response to Cu-withholding stress? How do genes encoded in the CRC respond to Cu-withholding stress? Thus far, only limited functional studies with *P. destructans* proteins have been reported^29–31^ and the tools to knock out genes or tag *P. destructans* proteins are limited^32, 33^. Therefore, we sought to take an unbiased quantitative proteomics approach to investigate how the *P. destructans* proteome responds to Cu-stress and to develop a global picture of the *P. destructans* proteome when cultured under laboratory conditions.

In this report, we characterize the *P. destructans* proteome under laboratory culture conditions using chemically defined growth media. The Cu-specific chemical chelator bathocuproine sulfonic acid (BCS) and copper sulfate were used to simulate Cu-withholding and Cu-overload growth conditions, respectively. While gross morphologic changes under Cu-stress conditions were not observed, the *P. destructans* proteomes were distinct, and several unique differentially abundant proteins (DAPs) were identified. Our studies indicate that chronic Cu-withholding stress results in greater alterations in the global *P. destructans* proteome than Cu-overload stress. These findings suggest that *P. destructans* must substantially adapt its proteome to overcome Cu restriction, while more modest adjustments are needed to withstand chronic Cu overload. Here, we also report the development and validation of a collection of antisera against *P. destructans* antigens that can be utilized to indicate Cu-withholding stress.

Together, this report provides an unbiased snapshot of the *P. destructans* proteome, along with antisera against *P. destructans* targets, which can be used to advance *P. destructans* molecular and metal cell biology.

## 2. Materials and Methods

### 2.1 Media preparation and propagation of Pseudogymnoascus destructans

The *Pseudogymnoascus destructans* strain (MYA-4855; ATCC strain number 20631-21) was purchased from the American Tissue Culture Collection (ATCC; Manassas, VA, USA) and used for all experiments. The long-term cultivation of *P. destructans* was carried out on solid yeast extract-peptone and glucose (YPD) plates at 15 °C, which contained 10 g/L of yeast extract, 20 g/L of peptone, 20 g/L of glucose, and 15 g/L of agar containing 50 μg/ml gentamycin. Metal growth studies were performed using chemically defined growth plates (SC-Ura) which were made using Milli-Q processed water and 1.8 g/L of yeast nitrogen base, 5 g/L ammonium sulfate (NH_4_SO_4_), 20 g/L glucose, 100 mg/L histidine, 1.7 g/L of the SC-Ura-His amino acid supplement mixture (Sunrise Scientific, Knoxville, TN), and 15 g/L agar. Before autoclaving, 1 pellet (∼200 mg) of NaOH was added to the SC-Ura media to assist in agar gel formation. Sterile Cu-sulfate, Fe-sulfate, and bathocuproine sulfonic acid sodium salt (BCS) were added to the SC-Ura growth media using a 125 mM stock solution after autoclave sterilization at approximately 50 °C. The control growth conditions consisted of SC-Ura plates supplemented with a final concentration of 10 μM Cu- and Fe-sulfate. Copper-withholding and copper-overload growth conditions were simulated in SC-Ura by the addition of the copper chelator BCS (final concentration 800 µM) or CuSO_4_ (final concentration 500 µM), respectively.

The inoculation of SC-Ura experimental growth plates was performed as follows: *Pseudogymnoascus destructans* conidia and mycelium were harvested from a working YPD plate, grown for two weeks at 15 °C, by the addition of 2 mL of TE buffer (10 mM Tris/1 mM disodium ethylenediaminetetraacetic acid (Na_2_-EDTA; pH = 7.4) to the plate and gently rubbing the mycelium mass with an inoculation loop. The resulting cell suspension was transferred and passed through a sterile 20-micron cell strainer to remove *Pd* hyphal cells. 100 µL of the resulting filtrate was used to inoculate experimental growth plates.

### 2.2 Generation of Pd CRC and PdCtr1b antiserum

Custom antiserum products were produced using the standard 90-New Zealand rabbit antibody service from Cocalico Biologicals, Inc. [Reamstown, PA, USA; Assurance number D16-00398 (A3669-01)]. The peptide and protein antigen targets are listed in Supplemental Table S1. For peptide-based antigens, two unique peptides were synthesized for each protein target and coupled to keyhole limpet hemocyanin (KLH) as the carrier protein. The co-injection of two peptide/KLH constructs was used to produce antibodies in rabbit hosts.

### 2.3 P. destructans protein extraction for proteomics analysis

*P. destructans* fungi were grown for 10 days at 15 °C on SC-Ura plates. Fungal cells, including conidia and mycelium, were suspended in 2 mL of TE buffer by gently rubbing the mycelium mass with a sterile inoculating loop. The resulting cell suspension was transferred to a 2-mL microcentrifuge tube and pelleted for 2 min at 16,000 X g. The cell pellet was washed twice by resuspending in 1 mL of TE buffer and pelleted by centrifugation. The resulting cell pellet was then snap-frozen in dry ice and stored at −80 °C for later use.

Total *P. destructans* cellular proteins were isolated using a modified trichloroacetic acid (TCA) precipitation protocol described by Anne et al^29^. Briefly, *P. destructans* cells were lysed using 300 μL of TCA buffer (10 mM Tris-HCl, 10% TCA, 25 mM ammonium acetate and 1 mM Na_2_-EDTA) with 100 μL of 0.5 μM zirconium beads (ZB-05; Next Advanced Inc., Troy, NY, USA) at Speed 12 for 5 min at 4 °C in a Bullet Blender 5E gold (Next Advanced Inc., Troy, NY, USA). The resulting total cellular lysate was transferred to a new tube, leaving the zirconium beads behind. The transferred material was centrifuged at 4 °C at 16,000 X g for 10 min, and the supernatant was discarded. The resulting pellet was frozen −80 °C and transferred to the University of Texas Health Science Center San Antonio Institutional Mass Spectrometry Core Laboratory.

Cells were lysed in buffer containing 5% SDS/50 mM triethylammonium bicarbonate (TEAB) in the presence of protease and phosphatase inhibitors (Halt; Thermo Scientific) and nuclease (Pierce™ Universal Nuclease for Cell Lysis; Thermo Scientific). After centrifugation, aliquots of the supernatants containing 85 µg of protein (EZQ™ Protein Quantitation Kit; Thermo Scientific) were mixed with a buffer containing 10% SDS/50 mM triethylammonium bicarbonate (TEAB), reduced with tris(2-carboxyethyl)phosphine hydrochloride (TCEP) and alkylated in the dark with iodoacetamide. After quenching with dithiothreitol, 12% phosphoric acid solution was added to each sample and the mixtures were applied to S-Traps (micro; Protifi) for tryptic digestion (sequencing grade; Promega) for 2 hr at 37 °C in 50 mM triethylammonium bicarbonate (TEAB). Peptides were eluted from the S-Traps sequentially with 50 mM TEAB, 0.2% formic acid, and 0.2% formic acid in 50% aqueous acetonitrile The pooled eluates were dried by vacuum centrifugation, redissolved in starting HPLC mobile phase (3% B, see below). and quantified using Pierce™ Quantitative Fluorometric Peptide Assay (Thermo Scientific).

Digests were analyzed by data-independent acquisition mass spectrometry on an Orbitrap Fusion™ Lumos™ Tribrid™ Mass Spectrometer (Thermo Fisher). On-line HPLC separation was accomplished with an RSLCnano HPLC system (Thermo Scientific/Dyonex): column, PicoFrit™ (75 μm internal diameter.; New Objective; Littleton, MA/USA) packed to 15 cm with C18 adsorbent (218MS 5 μm, 300 Å; Vydac/Grace; Columbia, MD/USA); mobile phase A, 0.5 % acetic acid (HAc)/0.005 % trifluoroacetic acid (TFA) in water; mobile phase B, 90 % acetonitrile/0.5 % HAc/0.005 % TFA/9.5 % water; gradient 3 – 42 % B in 120 min; flow rate, 0.4 μL/min. A pool was made of the experimental samples, and 2-µg peptide aliquots were analyzed using three stages of gas-phase fractionation (395–605 m/z, 595–805 m/z, 795–1005 m/z, staggered) and 4-m/z windows (30k resolution for precursor and product ion scans, all in the orbitrap). The resulting three data files were used to create an empirically-corrected DIA chromatogram library^34^ by searching against a Prosit-generated predicted spectral library^35^ based on a full UniProt *Pseudogymnoascus destructans* (Strain ATCC MYA-4855 / 20631-21) protein sequence database (UP000011064_658429 downloaded on 20240812). Experimental samples were randomized for sample preparation and analysis; injections of 2 µg of peptides and a 2-hr HPLC gradient were employed. MS data for experimental samples were acquired in the orbitrap using 8-m/z windows (staggered; 30k resolution for precursor and product ion scans) and searched against the chromatogram library. Scaffold DIA (v3.4.1; Proteome Software) was used for all DIA-MS data processing: fixed modification, cysteine carbamidomethylation; proteolytic enzyme, trypsin with one missed cleavage allowed; peptide mass tolerance, ±10.0 ppm; fragment mass tolerance, ±10.0 ppm; charge states, 2+ and 3+; peptide length, 6–30. Peptides identified in each sample were filtered by Percolator^36^ to achieve a maximum FDR of 1%. Individual search results for each sample type were combined and peptide identifications were assigned posterior error probabilities and filtered to an FDR threshold of 1% by Percolator^36^. Peptide quantification was performed by Encyclopedia^34^ based on the three to five highest quality fragment ions. Only peptides that were exclusively assigned to a protein were used for relative quantification unless specified otherwise.

### 2.4 Western blot analysis

*P. destructans* was cultured with conidia and mycelium isolated exactly as described in 2.3. Total cellular proteins were collected using a trichloroacetic acid (TCA) extraction protocol as previously described^29^ with total protein levels estimated using a BCA Protein Assay Kit (Pierce Thermofisher Waltham, MA, USA) according to the manufacturer’s specifications. SDS-PAGE and western blot analysis were performed using 30 µg/lane of total protein. Before transferring the protein to a PVDF membrane, the gels were stained with 4% trichloroethanol (TCE) solution for an hour and then washed with deionized water ^37, 38^. The gels were imaged using the Bio-Rad stain-free gel system to assess loading levels. For antigen detection, the PVDF membrane was incubated with TSU antiserum (see Supplemental Table S1 for dilution ratio) in 1X TBST containing 5% non-fat milk powder for an hour. After incubation, the membrane was washed three times with 1X TBST for 5 minutes each. Finally, the membrane was incubated with a goat anti-rabbit CY3 (Cell Signaling Technology, Danvers, MA, USA) secondary antibody at a 1:5,000 dilution for thirty minutes. Excess secondary antibody was removed by washing the membrane three times with 1X TBST for 5 minutes each. Western blot images were collected using a Bio-Rad ChemiDoc MP imaging system (Bio-Rad Laboratories, Inc, Hercules, CA, USA).

### 2.5 Pseudogymnoascus destructans microscopy

The *P. destuctans* fungus was grown on SC-Ura plates as described in 2.3 and inoculated with an initial seed density of 250,000 cells per plate. Harvested cells from experimental growth plates were passed through a 20-micron cell strainer to remove *Pd* hyphal cells. The resulting spores were pelleted at 5000 x g for five minutes and the supernatant was removed. The resulting pellet was then resuspended in 200 μL of Karnovsky’s fixative (20 mL of 16% paraformaldehyde solution, 8 mL of 50% glutaraldehyde EM grade, 25 mL of 0.2M sodium phosphate buffer, and 25 mL of distilled water; Electron Microscopy Sciences, Hatfield, PA USA). The solution was incubated at room temperature for five minutes and then incubated at 4 °C overnight. Following incubation, the sample was centrifuged at 3,000 x g at 4 °C for ten minutes and the supernatant was discarded. The pellet was resuspended in equal volume 50 mM Calcoflour White for 10 minutes at room temperature. The sample was centrifuged at 3,000 x g at 4 °C for 10 minutes and then washed with 1X PBS two times to remove excess Calcoflour White. After washing, the pellet was resuspended in 100 μL of 1X PBS. Five μL of the stained cell solution and 5 μL of glycerol were added to a slide. The cover slip was added and sealed with clear nail polish. Microscope images were collected on a Zeiss Axio Observer 7 system at 1000X total magnification equipped with an 820 monochrome digital camera, aptotome-3, and DAPI/GFP filter sets. The Zeiss Zen software (version 3.8) package was used for image processing.

### 2.6 Bioinformatic Methods

Scaffold DIA (v3.4.1; Proteome Software; Portland, OR, USA) was used to generate pairwise comparison tables using a pairwise t-test versus control samples with a significance threshold value of 0.05. Reported fold changes (FC) are calculated based on exclusive intensity versus control growth conditions unless specified otherwise. UniProt identifiers were mapped to VC83_gene identifiers using an in-house BLAST comparison. GO terms assigned based on the NCBI refseq VC83 gene annotation (ASM1612v1; assessed 09/2025). Unless otherwise specified, proteins were considered significantly abundant for Cu-withholding growth conditions when the |Log_2_FC| > 1.5, *p* < 0.05 compared to control growth conditions. Whereas, proteins were considered significantly abundant under Cu-overload growth conditions when |Log_2_FC| > 1.0, *p* < 0.05.

## 3. Results

### 3.1 P. destructans growth behavior and spore morphology under chronic Cu-stress growth conditions

*Pseudogymnoascus destructans* cultured on SC-Ura growth media under extremes of Cu bioavailability have demonstrated that *P. destructans* displays robust growth behavior with minimal impacts on its characteristic asymmetric curved conidia morphology^39^. In this study, we supplemented SC-Ura control growth conditions with 10 μM Cu-sulfate and Fe-sulfate to bolster metal availability during large-scale cultivation conditions needed for protein isolation. Under our growth conditions, we observed a distinct grey-green hyphal mass on the top of the growth plates of the control and Cu-overload growth conditions, whereas *P. destructans* cultured on the Cu-withholding (i.e., SC-Ura + 800 μM BCS) growth plates display a distinct white hyphal mass (Figure 1). Spores isolated from the three growth conditions used in this study displayed similar curved conidia morphologies under brightfield microscopy at 10,000 x total magnification with a mean length of 4.6 (± 0.6) microns (n = 10) and width of 1.9 (± 0.2) microns (n = 10) (Supplemental Figure S1).

**Figure 1.**
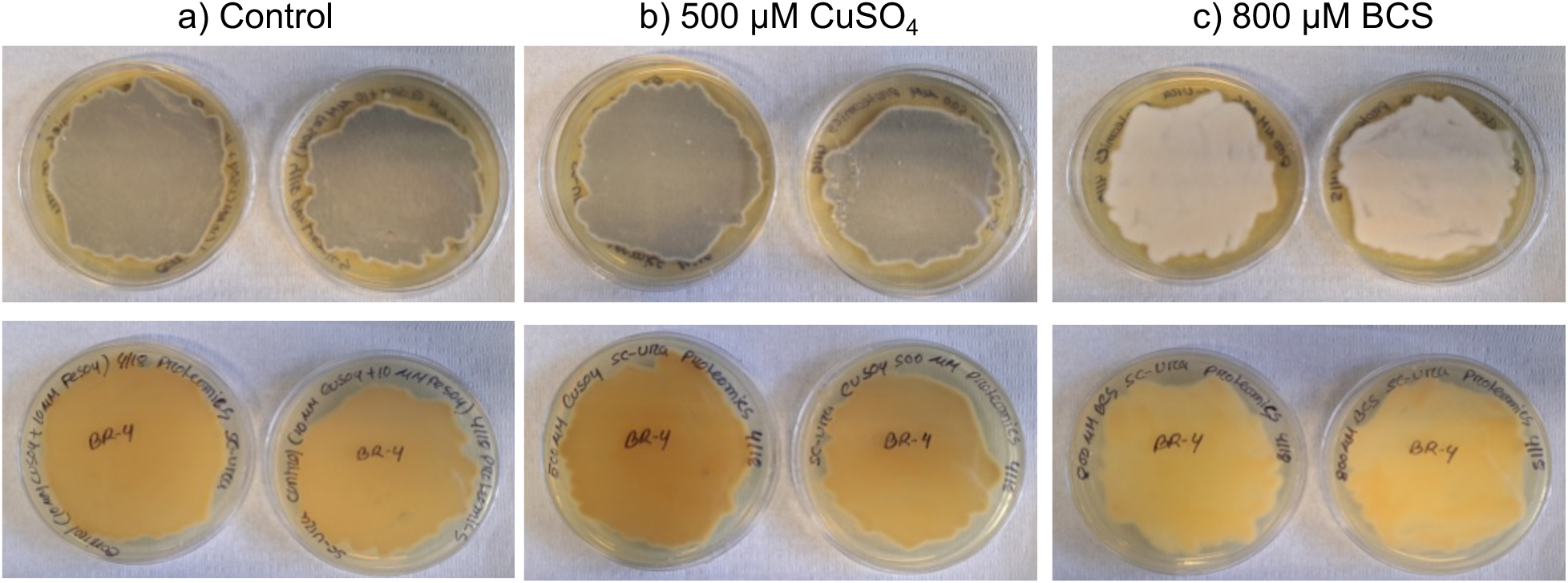
Culture plates of *P. destructans* under varying Cu-stress growth conditions. *P. destructans* on growth plates consisting of (a) SC-Ura +10 μM CuSO_4_ and +10 μM FeSO_4_ (control), (b) SC-Ura + 500 μM CuSO_4_ (Cu-overload), (c) SC-Ura + 800 μM BCS (Cu-withholding).

### 3.2 P. destructans global proteomic profiles

Across all experimental conditions tested, data-independent acquisition mass spectrometry (DIA-MS) identified 4340 proteins (47.8%) of the total 9073 predicted *P. destructans* proteome (Supplemental Table S2). The dynamic intensity range of detected proteins spans 6.1 orders of magnitude (Figure 2a; Supplemental Table S2). Principal component analysis (PCA) plots of control and Cu-stress samples show a large separation between the Cu-withholding and control/Cu-overload data sets (Figure 2b, red versus blue or green). A closer clustering pattern is observed for the Cu-overload and control data sets (Figure 2b, blue versus green).

**Figure 2.**
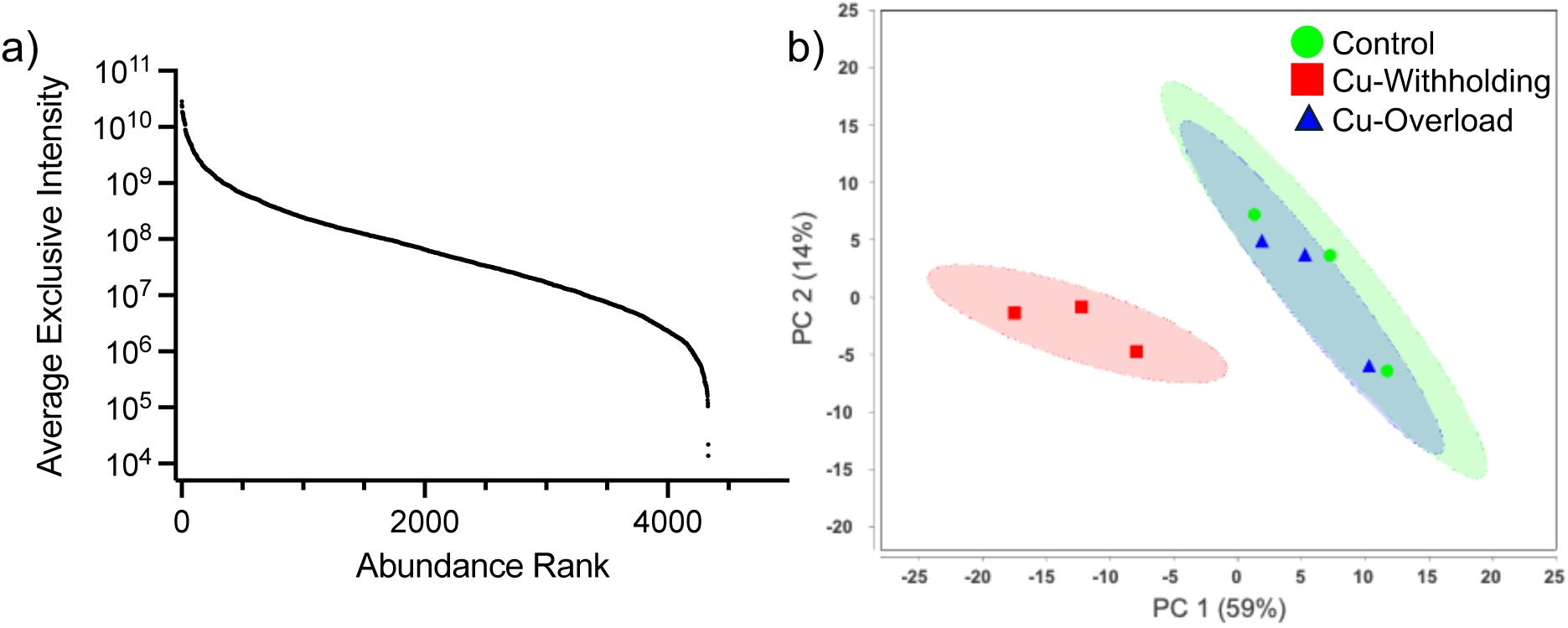
Characterization of the *P. destructans* proteome under control and Cu-stress growth conditions. (a) Average exclusive peptide intensity levels for proteins identified in whole-cell *P. destructans* lysates. Proteins are organized by relative intensity rank and plotted on a log_10_ scale to accommodate plotting all proteins. (B) Principal component analysis plot comparing global protein levels in control (green), Cu-withholding (red), and Cu-overload (blue) conditions. Colored ellipses represent the 95% confidence levels for each sample.

Analysis of the Cu-withholding treated cells versus control cells identifies a total of 1395 differentially abundant proteins (DAPs) (*i.e.,* 32% of the detectible proteome) displaying statistically significant p < 0.05 changes in protein relative abundance (Supplemental Table S3). This includes 590 DAPs that show higher abundance and 805 DAPs that are less abundant than in cells grown under control conditions. In comparison, a total of 387 DAPs (i.e., 9.0 % of the detectible proteome) display statistically significant (p < 0.05) changes in abundance levels in the Cu-overload treatment versus control conditions. Copper overload stress leads to 200 DAPs that are less abundant and 187 DAPs that are more abundant vs control growth conditions (Supplemental Table S4). Due to the large number of DAPs identified in the pairwise analysis, a more stringent fold-change (FC) threshold was applied for further analysis. From here on, we performed global analysis on DAPs under Cu-withholding (BCS) stress displaying a |Log_2_FC| > 1.5 (i.e., 2.8x), *p* < 0.05, and under Cu-overload stress displaying a |Log_2_FC| > 1.0 (i.e., 2x), *p* < 0.05, when compared to control cells. Volcano plots of the results for Cu-withholding and Cu-overload stress are displayed in Figure 3.

**Figure 3.**
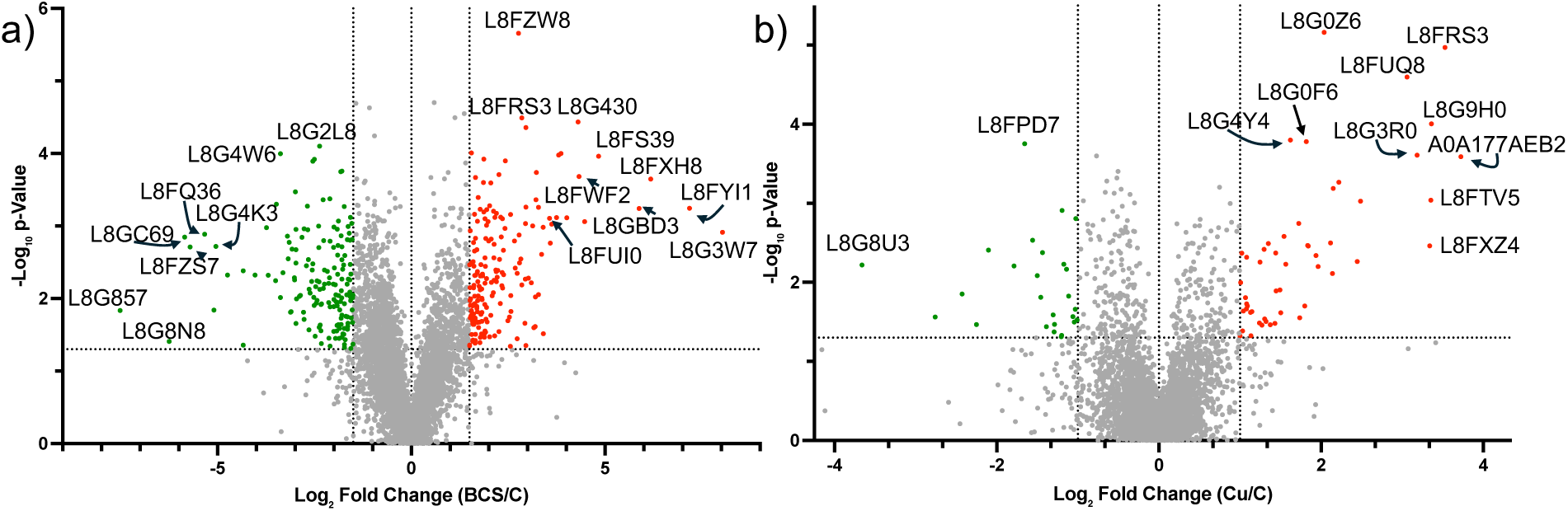
Volcano plots of global proteomic analysis upon Cu-withholding and Cu-overload stress. (a) Cu-withholding (BCS) vs control. DAPs that display a |log_2_FC| > 1.5, FDR *p* < 0.05 threshold are colored in red (increased abundance) and green (decreased abundance). Notable proteins are identified. (b) Cu-overload (Cu) and control samples. DAPs that display a |log_2_FC| > 1.0, FDR *p* < 0.05 threshold are indicated in red (increased abundance) and green (decreased abundance). Notable proteins are identified.

Employing a higher fold-change qualifier, we identify 343 Cu-withholding and 78 Cu-overload DAPs when compared to control growth conditions. These DAPs are widely distributed over the range of detectable proteins (Figure 4a red, blue, and purple dots). There are 44 DAPs that are common to both Cu-withholding and Cu-overload stress conditions, 299 DAPs are unique to Cu-withholding stress, and 34 DAPs are unique to Cu-overload stress (Figure 4b, Supplemental Table S5).

**Figure 4.**
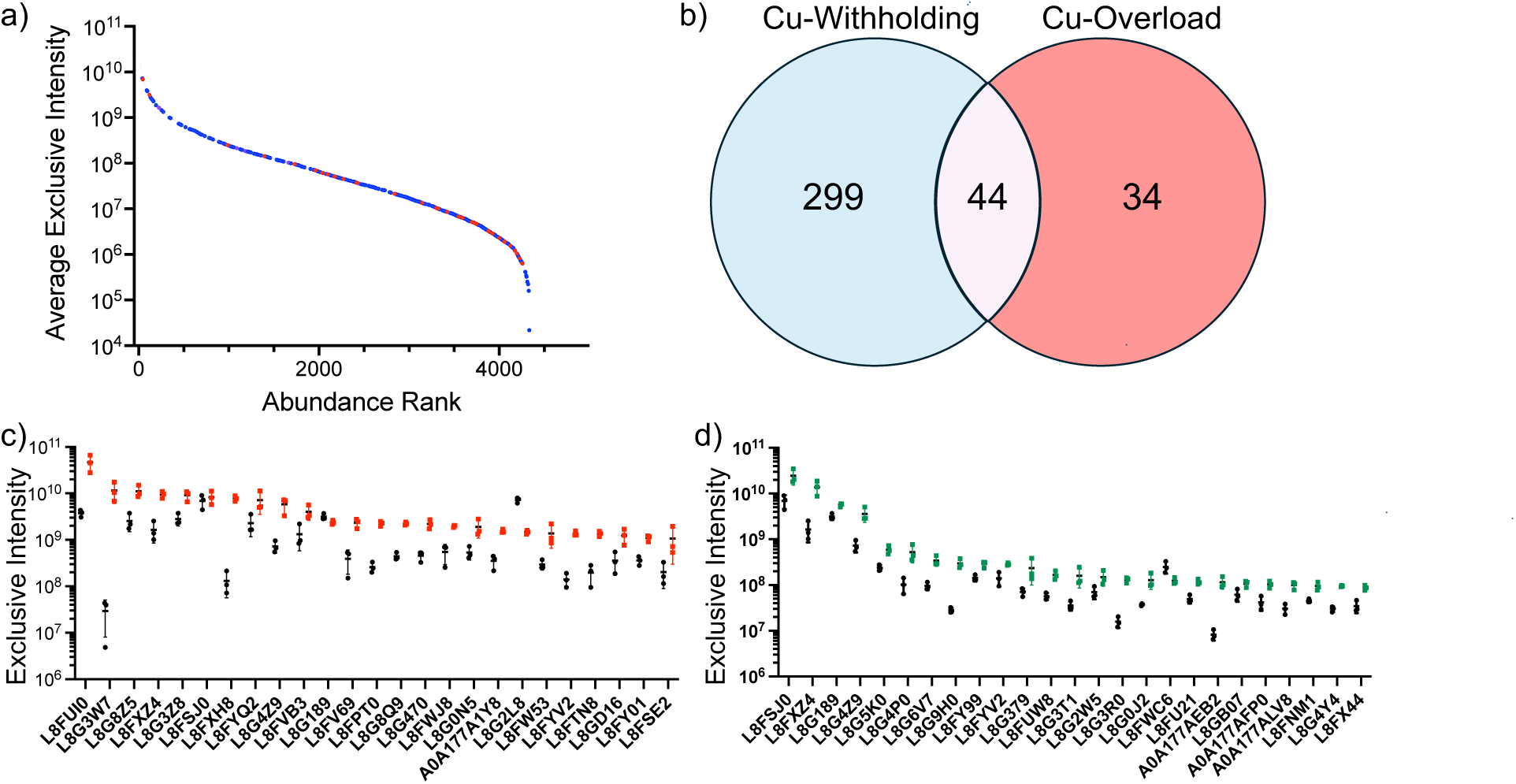
Analysis of identified DAPs found under Cu-withholding and Cu-overload growth conditions. (a) Intensity plot identifying DAPs indicated by rank under control growth conditions. DAPs unique to Cu-withholding (blue) or Cu-overload (red). DAPs common to both Cu-withholding and Cu-overload stress are colored purple. (b) Venn diagram of DAPs identified under Cu-withholding or Cu-overload growth conditions. A total of 229 DAPs are unique to Cu-withholding samples, 34 DAPs are unique to Cu-overload samples, and 44 DAPs are common to both Cu-withholding and Cu-overload conditions. (c) Plot of the 25 most abundant DAPs identified under Cu-withholding conditions. Protein levels in Cu-withholding samples are indicated in red (n = 3) and control samples are indicated in black (n = 3). (d) Plot of the 25 most abundant DAPs identified under Cu-overload conditions. Protein levels in control samples are displayed in black (n = 3) and Cu-overload samples are displayed in green (n = 3). For plots in panels (c) and (d), each point represents a biological replicate with the mean and standard deviation of the exclusive intensity indicated by horizontal and vertical lines, respectively.

We further identified the 25 most abundantly detected DAPs based on total exclusive intensity (Table 1 and Table 2) and the 25 DAPs displaying the largest fold changes (Table 3 and Table 4). The former may be used as a proxy to identify DAPs with high sensitivity to detection. Based on our analysis, it appears that chronic Cu-withholding stress has a more severe impact on the global *P. destructans* proteome than chronic Cu-overload stress. This is based on the number of DAPs and the magnitude of relative changes vs control growth conditions. The average change for DAPs under Cu-withholding stress |Log_2_FC| = 2.35, whereas the average |Log_2_FC| of DAPs under Cu-overload stress is 1.69 (Supplemental Table S5). Furthermore, only 19 DAPs display large |Log_2_FC| values > 2 under Cu-overload conditions. These changes are reflected in the relative detection levels of the 25 most abundantly detected DAPs under Cu-withholding and Cu-overload stress displayed in Figures 4c and 4d, respectively. We find that the 25 most abundant detected DAPs (based on total exclusive intensity) under Cu-withholding stress are highly represented in the most abundant globally detected proteins found in Cu-starved cells (Figure 4c and Table 1); all 25 DAPs are found in the top 375 of globally detected proteins with seven DAPs represented in the most abundant 50 proteins. In contrast, the 25 most abundant DAPs identified in chronic Cu-overload stressed cells (based on total exclusive intensity counts) are more widely distributed across the detectable *P. destructans* proteome, and only three DAPs were identified within the most abundant 100 detected proteins (Figure 4d and Table 2). Surprisingly, many DAPs which display the largest Log_2_FC increases (Table 3 and Table 4) also readily detectable in *P. destructans* samples (Table 1 and Table 2).

**Table 1.**
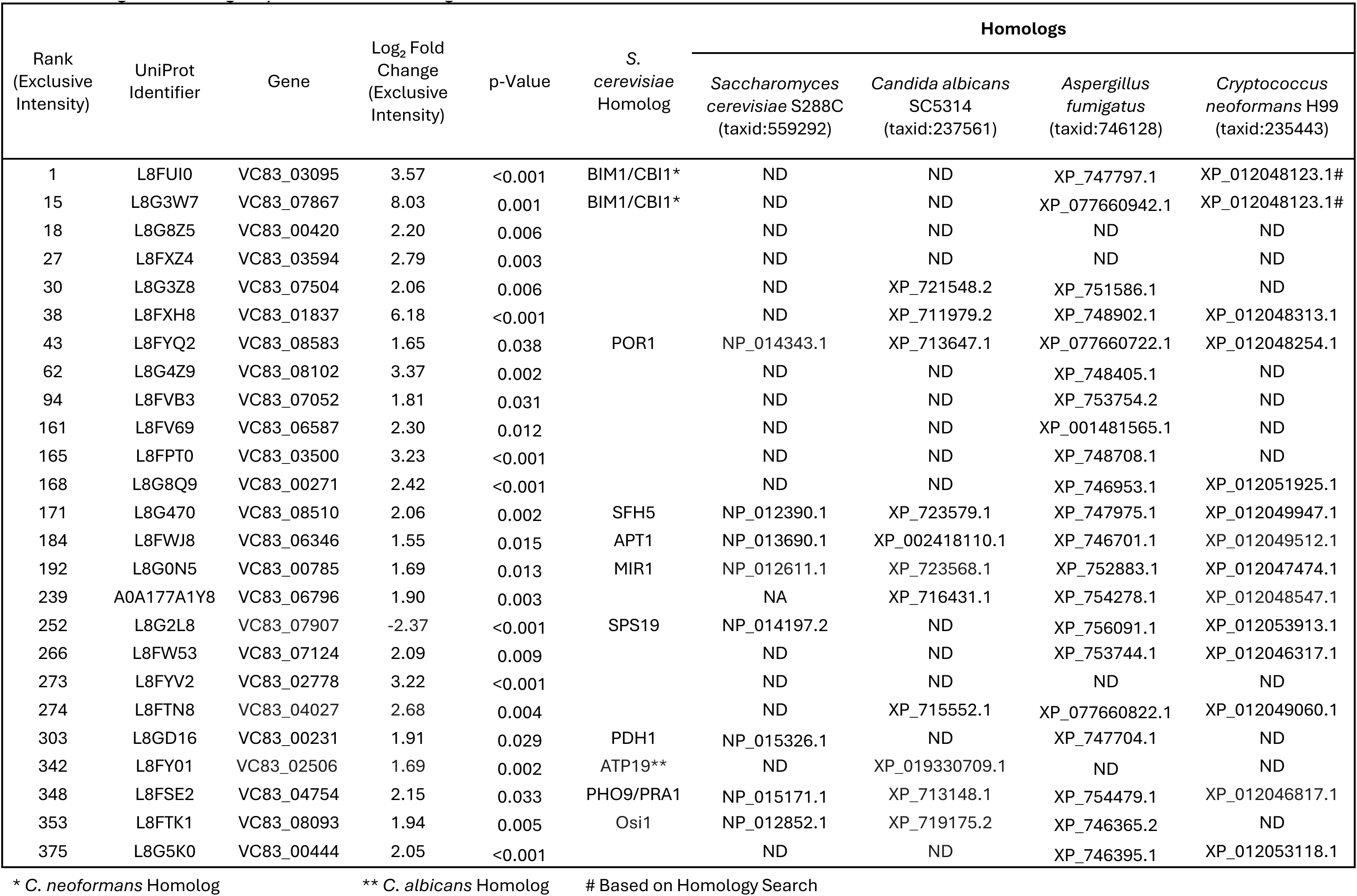
The 25 most abundant DAPs based on MS intensity and their respective fungal homologs upon Cu-withholding stress.

**Table 2.** The 25 most abundant DAPs based on MS intensity and their respective fungal homologs upon Cu-overload stress.

| Rank<br>(Exclusive<br>Intensity) | UniProt<br>Identifier | Gene | Log<sub>2</sub> Fold<br>Change<br>(Exclusive<br>Intensity) | p-Value | S.<br>*cerevisiae*<br>Homolog | Homologs |  |  |  |
| --- | --- | --- | --- | --- | --- | --- | --- | --- | --- |
|  |  |  |  |  |  | *Saccharomyces*<br>*cerevisiae* S288C<br>(taxid:559292) | *Candida*<br>*albicans* SC5314<br>(taxid:237561) | *Aspergillus*<br>*fumigatus*<br>(taxid:746128) | *Cryptococcus*<br>*neoformans* H99<br>(taxid:235443) |
| 5 | L8FSJ0 | VC83_09026 | 1.49 | 0.012 |  | ND | ND | ND | ND |
| 17 | L8FXZ4 | VC83_03594 | 3.34 | 0.003 |  | ND | ND | ND | ND |
| 73 | L8G189 | VC83_06745 | 1.02 | 0.004 |  | ND | ND | XP_748486.1 | XP_012050966.1 |
| 113 | L8G4Z9 | VC83_08102 | 2.12 | 0.003 |  | ND | ND | XP_748405.1 | ND |
| 547 | L8G5K0 | VC83_00444 | 1.30 | 0.004 |  | ND | ND | XP_746395.1 | XP_012053118.1 |
| 599 | L8G4P0 | VC83_08083 | 2.14 | 0.008 |  | ND | ND | ND | ND |
| 811 | L8G6V7 | VC83_01748 | 1.72 | 0.002 | ASM3** | NP_010740.3 | XP_718999.1 | XP_752162.1 | XP_012053233.1 |
| 887 | L8G9H0 | VC83_05770 | 3.36 | <0.001 |  | ND | ND | ND | ND |
| 909 | L8FY99 | VC83_08999 | 1.25 | 0.006 |  | ND | ND | ND | XP_012049737.1 |
| 913 | L8FYV2 | VC83_02778 | 1.04 | 0.023 |  | ND | ND | ND | ND |
| 1040 | L8G379 | VC83_04907 | 1.30 | 0.029 |  | ND | ND | XP_001481417.1 | ND |
| 1263 | L8FUW8 | VC83_08087 | 1.54 | 0.003 |  | ND | ND | XP_747896.1 | ND |
| 1291 | L8G3T1 | VC83_01092 | 1.96 | 0.006 | FET3 | NP_013774.1 | XP_711267.2 | XP_747965.2 | XP_012053059.1 |
| 1355 | L8G2W5 | VC83_02399 | 1.13 | 0.047 | RPS27A | NP_012766.1 | XP_002422270.1 | XP_754942.1 | XP_012051162.1 |
| 1461 | L8G3R0 | VC83_08103 | 3.18 | <0.001 |  | ND | ND | XP_756138.1 | ND |
| 1480 | L8G0J2 | VC83_07150 | 1.44 | 0.004 | FET3 | NP_013774.1 | XP_711267.2 | XP_747965.2 | ND |
| 1518 | L8FWC6 | VC83_05646 | -1.03 | 0.022 | COX4 | NP_011328.1 | XP_722521.1 | XP_749436.2 | XP_012048730.1 |
| 1554 | L8FU21 | VC83_00654 | 1.35 | 0.003 | ETR1 | NP_009582.1 | XP_713016.1 | XP_748653.1 | ND |
| 1572 | A0A177AEB2 | VC83_03593 | 3.72 | <0.001 |  | ND | ND | ND | ND |
| 1628 | L8GB07 | VC83_06840 | 1.03 | 0.041 | URE2 | NP_014170.1 | XP_720446.1 | XP_748959.1 | XP_012051409.1 |
| 1657 | A0A177AFP0 | VC83_02698 | 1.44 | 0.013 | VRP1 | NP_013441.1 | XP_716030.2 | ND | XP_012049812.1 |
| 1693 | A0A177ALV8 | VC83_00142 | 1.84 | 0.003 |  | ND | ND | XP_077661402.1 | ND |
| 1738 | L8FNM1 | VC83_08526 | 1.08 | 0.005 |  | ND | ND | XP_077661321.1 | ND |
| 1740 | L8G4Y4 | VC83_08076 | 1.62 | <0.001 |  | ND | ND | XP_750704.2 | ND |
| 1798 | L8FX44 | VC83_08099 | 1.56 | 0.006 | ERG26 | NP_011514.1 | XP_715564.1 | XP_756075.1 | ND |

\*\* *C. albicans* Homolog

**Table 3.**
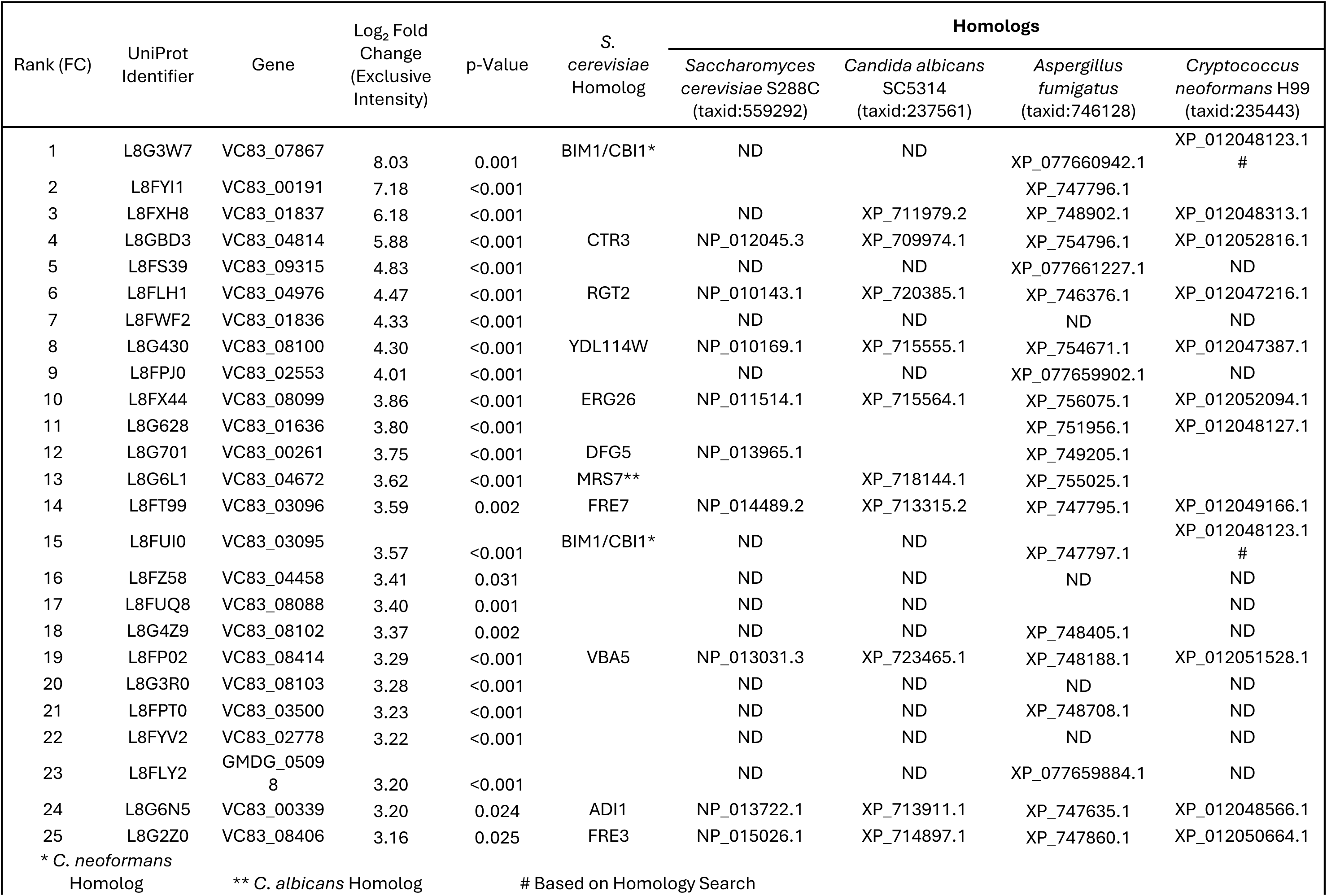
Table of the 25 DAPs displaying largest increases upon Cu-withholding stress.

**Table 4.** Table of the 25 DAPs displaying highest increases upon Cu-overload stress.

| Rank<br>(FC) | UniProt<br>Identifier | Gene | Log<sub>2</sub> Fold<br>Change<br>(Exclusive<br>Intensity) | p-Value | S.<br>*cerevisiae*<br>Homolog | Homologs |  |  |  |
| --- | --- | --- | --- | --- | --- | --- | --- | --- | --- |
|  |  |  |  |  |  | *Saccharomyces*<br>*cerevisiae* S288C<br>(taxid:559292) | *Candida albicans*<br>SC5314<br>(taxid:237561) | *Aspergillus*<br>*fumigatus*<br>(taxid:746128) | *Cryptococcus*<br>*neoformans* H99<br>(taxid:235443) |
| 1 | A0A177AEB2 | VC83_03593 | 3.72 | <0.001 |  | ND | ND | ND | ND |
| 2 | L8FRS3 | VC83_09319 | 3.53 | <0.001 |  | ND | ND | XP_749913.1 | ND |
| 3 | L8G9H0 | VC83_05770 | 3.36 | <0.001 |  | ND | ND | ND | ND |
| 4 | L8FTV5 | VC83_00653 | 3.35 | <0.001 |  | ND | ND | XP_754100.1 | ND |
| 5 | L8FXZ4 | VC83_03594 | 3.34 | 0.003 |  | ND | ND | ND | ND |
| 6 | L8G3R0 | VC83_08103 | 3.18 | <0.001 |  | ND | ND | ND | ND |
| 7 | L8FUQ8 | VC83_08088 | 3.06 | <0.001 |  | ND | ND | ND | ND |
| 8 | L8FW89 | VC83_07721 | 2.49 | <0.001 | CCC2 | NP_010556.1 | XP_723470.2 | XP_754347.1 | XP_012053482.1 |
| 9 | L8FLH1 | VC83_04976 | 2.44 | 0.005 | RGT2 | NP_010143.1 | XP_720385.1 | XP_746376.1 | XP_012047216.1 |
| 10 | L8FPW3 | VC83_01431 | 2.22 | <0.001 | ALD3 | NP_013892.1 | XP_721543.1 | XP_747354.1 | XP_012050896.1 |
| 11 | L8G430 | VC83_08100 | 2.15 | <0.001 | YDL114W | NP_010169.1 | XP_715555.1 | XP_754671.1 | XP_012047387.1 |
| 12 | L8G4P0 | VC83_08083 | 2.14 | 0.008 |  | ND | ND | ND | ND |
| 13 | L8G4Z9 | VC83_08102 | 2.12 | 0.003 |  | ND | ND | XP_748405.1 | ND |
| 14 | L8G0Z6 | VC83_08082 | 2.04 | <0.001 |  | ND | ND | XP_754149.1 | ND |
| 15 | L8G3T1 | VC83_01092 | 1.96 | 0.006 | FET3 | NP_013774.1 | XP_711267.2 | XP_747965.2 | XP_012053059.1 |
| 16 | L8G085 | VC83_08518 | 1.94 | 0.005 | JLP1 | NP_013043.1 | XP_720050.1 | XP_750960.1 | XP_012050939.1 |
| 17 | A0A177ALV8 | VC83_00142 | 1.84 | 0.003 |  | ND | ND | XP_077661402.1 | XP_012047221.1 |
| 18 | L8FS39 | VC83_09315 | 1.83 | 0.003 |  | ND | ND | XP_077661227.1 | ND |
| 19 | L8G0F6 | VC83_05017 | 1.82 | <0.001 |  | ND | ND | XP_749118.2 | ND |
| 20 | L8G5U3 | VC83_08040 | 1.80 | 0.020 |  | ND | ND | ND | ND |
| 21 | L8G9Q9 | GMDG_03454 | 1.74 | 0.028 |  | ND | ND | ND | ND |
| 22 | L8G6V7 | VC83_01748 | 1.72 | 0.002 | ASM3* | NP_010740.3 | XP_718999.1 | XP_752162.1 | XP_012053233.1 |
| 23 | L8G4Y4 | VC83_08076 | 1.62 | <0.001 | DIT2 | NP_010690.1 | ND | XP_750704.2 | ND |
| 24 | L8FX44 | VC83_08099 | 1.56 | 0.006 | ERG26 | NP_011514.1 | XP_715564.1 | XP_756075.1 | XP_012052094.1 |
| 25 | L8FUW8 | VC83_08087 | 1.54 | 0.003 |  | ND | ND | XP_747896.1 | ND |

\* *C. albicans* Homolog

### 3.3 P. destructans proteome under chronic Cu-withholding stress

This study aimed to understand how *P. destructans* adapts its proteome in response to Cu-withholding stress and to understand if the adaptations observed at the transcriptional level are mirrored at the protein level. Our analysis focused primarily on adaptations related to cellular redox homeostasis or mitochondrial function, superoxide dismutase (SOD) enzymes, and enzymes associated with the cell surface and extracellular environment which have been previously implicated in the *P. destructans* Cu-stress response^23, 29^.

#### 3.3.1 Reorganization of the mitochondrial electron transport chain

As in all eukaryotes, mitochondria play an important role in maintaining energy flux by coupling carbon metabolism to respiration. Within the top 25 most abundant detected DAPs impacted by Cu-withholding stress (Table 1), we identify three mitochondrial proteins involved in maintaining redox and energy homeostasis. Specifically, the proteins L8FYQ2 (a POR1 homolog), L8G0N5 (a PIC2/MIR1 homolog), and L8FY01 (an ATP19A homolog) all display an approximate three-fold increase in protein levels upon Cu-restriction. Together, these enzymes assist in maintaining redox homeostasis through mitochondrial glutathione trafficking,^40^ mitochondrial Cu-import,^41^ and ATP synthase activity^42^. Given the high representation of mitochondrial proteins among the top 25 most abundant detected DAPs under Cu-withholding stress, and copper’s role as an essential redox cofactor in cytochrome *c* oxidase (CcO), we examined changes in annotated CcO subunits, CcO assembly proteins, and alternative oxidase (AOX) protein levels (Table 5).

**Table 5.** Regulation of genes associated with cytochrome *c* oxidase activity/assembly and respiration.

| UniProt Identifier | Gene | S. cerevisiae Homolog | Homologs |  |  |  | P-Value Cu vs C | P-Value BCS vs C | Log<sub>2</sub>FC Cu vs C | Log<sub>2</sub>FC BCS vs C |
| --- | --- | --- | --- | --- | --- | --- | --- | --- | --- | --- |
|  |  |  | *Saccharomyces cerevisiae* S288C (taxid:559292) | *Candida albicans* SC5314 (taxid:237561) | *Aspergillus fumigatus* (taxid:746128) | *Cryptococcus neoformans* H99 (taxid:235443) |  |  |  |  |
| L8G6R7 | VC83_00757 | COX5 | NP_014346.1 | XP_723445.1 | XP_753588.1 | XP_012052894.1 | 0.176 | 0.092 | 0.45 | -0.44 |
| L8FRX7 | VC83_02346 | COX13 | NP_011324.1 | XP_722517.1 | XP_754987.2 | XP_012048235.1 | 0.938 | 0.005 | 0.10 | -3.02 |
| L8FPI1 | VC83_02644 | COX19 | NP_013082.1 | XP_019330723.1 | XP_001481718.1 | XP_012052175.1 | 0.120 | 0.206 | 0.60 | -0.56 |
| L8G8K5 | VC83_03145 | PET117<sup>#</sup> | ND | ND | XP_746730.1 | ND | 0.884 | 0.044 | 0.02 | 1.50 |
| L8G1A6 | VC83_03780 | COX12 | NP_013139.1 | XP_019331016.1 | XP_077660340.1 | XP_012050388.1 | 0.610 | 0.175 | -0.30 | -1.38 |
| L8FQD8 | VC83_04482 | COA3 | NP_076894.3 | ND | ND | XP_012046743.1 | 0.349 | <0.001 | -0.14 | 2.02 |
| L8G8A3 | VC83_04540 | COA6 | NP_013972.1 | XP_719365.1 | XP_754641.2 | XP_012048211.1 | 0.396 | 0.493 | 0.26 | -0.17 |
| L8FRT4 | VC83_04823 | AOX2<sup>*</sup> | ND | XP_723459.2 | XP_749637.1 | XP_012046575.1 | 0.111 | 0.001 | 0.60 | 2.78 |
| L8FZW8 | VC83_04895 | COX16 | NP_012531.1 | ND | XP_077660702.1 | XP_012051858.1 | 0.107 | <0.001 | -0.22 | 2.77 |
| L8FML5 | VC83_04970 | PET191 | NP_012568.1 | XP_019330823.1 | XP_001481507.1 | XP_012048575.1 | 0.240 | 0.366 | 0.38 | 0.59 |
| L8FWC6 | VC83_05646 | COX4 | NP_011328.1 | XP_722521.1 | XP_749436.2 | XP_012048730.1 | 0.022 | 0.013 | -1.03 | -1.39 |
| L8G0Y1 | VC83_07696 | COX15 | NP_011068.1 | XP_711328.2 | XP_750658.1 | XP_012053108.1 | 0.267 | 0.005 | -0.18 | 3.02 |
| L8G5Q2 | VC83_08913 | COX11 | NP_015193.1 | XP_711544.1 | XP_077660331.1 | XP_012047604.1 | 0.445 | 0.633 | 0.49 | 0.39 |
| L8FPX1 | VC83_08949 | COX6 | NP_011918.1 | XP_019330760.1 | XP_748067.1 | XP_012047506.1 | 0.722 | 0.106 | -0.06 | -1.61 |

\* homolog in *C. albicans*, <sup>#</sup>Superfamily identified through BLAST

In general, Cu-restriction causes large and significant changes in CcO assembly (COA) and CcO subunit protein levels. Increased levels of COA3 (L8FQD8, Log_2_FC = 2.02) and COX16 (L8FZW8, Log_2_FC = 2.77) homologs suggest that CcO assembly and COX1 subunit maturation may be impacted by copper restriction^43, 44^. Additionally, we also find a significant reduction in some CcO subunit proteins. Specifically, we observe an approximal 8- and 2.6-fold reduction in L8G6R7 (a COX5 homolog) and L8FWC6 (a COX4 homolog) CcO subunit levels, respectively. To our surprise, enzymes involved in CcO heme-cofactor assembly are also impacted upon Cu-withholding stress. The proteins L8G0Y1 (a COX15 homolog) and L8G8K5 (a PET117 homolog) which are responsible for heme-a biosynthesis and inserting of heme-a into CcO, also display significant increases under Cu-retriction^45, 46^. Taken together there is strong evidence that Cu-restriction highly impacts *P. destructans* CcO enzyme subunit levels and possible function.

Fungal pathogens such as *Candida albicans* can utilize an alternative oxidase (AOX) for respiration to bypass CcO as the site of terminal-O_2_ reduction when Cu levels are limited. AOX enzymes utilize a di-iron active site to achieve O_2_ reduction, whereas CcO has a high Cu-requirement—three Cu-ions per functional unit—to achieve efficient O_2_ reduction chemistry^16, 47^. The *P. destructans* genome encodes for one AOX enzyme, L8FRT4, for which we observe a 6.8-fold (Log_2_FC = 2.78) increase in protein levels in Cu-starved cells. The combination of large changes for several CcO chaperone and subunit protein levels in addition to increased levels of AOX protein suggests that extensive reorganization of the *P. destructans* mitochondrial electron transport chain occurs under Cu-withholding stress.

#### 3.3.2 Alteration in superoxide dismutase (SOD) levels and other Cu-responsive cytosolic enzymes

Previously, we identified a collection of genes, including the family of Cu- and Mn-containing superoxide dismutase (SOD) enzymes, as well as the Cu-responsive gene cluster (CRC), which display transcriptional profiles similar to those of *P. destructans* high-affinity Cu transporters *Pd*CTR1a and *Pd*CTR1b (*vida infra*)^23^. However, due to limited molecular probes for *P. destructans* proteins, we could not validate SOD and CRC expression levels^23^. In this current study, we observe similar Cu-regulatory patterns in protein levels for SOD, CRC, and other differentially expressed gene targets identified through transcriptional studies (Table 6) ^23^.

**Table 6.** Table of SOD proteins and previously identified genes regulated under Cu-stress conditions.

| UniProt Identifier | Gene | *P. destructans* gene name | P-Value Cu vs C | P-Value BCS vs C | Log<sub>2</sub>FC Cu vs C | Log<sub>2</sub>FC BCS vs C |
| --- | --- | --- | --- | --- | --- | --- |
| L8FTN0 | VC83_07077 | SOD1 | 0.076 | 0.009 | 0.96 | -1.57 |
| L8FYM0 | VC83_08495 | SOD4 | 0.190 | 0.500 | 0.99 | 0.48 |
| L8G9Z6 | VC83_07362 | SOD2 | 0.705 | 0.035 | 0.00 | 0.64 |
| L8G5D8 | VC83_02616 | SOD3 | 0.540 | 0.058 | -0.18 | 0.92 |
| L8FYI1 | VC83_00191 | *PdCTR1a* | 0.390 | <0.001 | -1.49 | 7.18 |
| L8GBD3 | VC83_04814 | *PdCTR1b* | 0.333 | <0.001 | -0.86 | 5.88 |
| L8FUI0 | VC83_03095 | *PdBLP2* | 0.287 | <0.001 | -0.56 | 3.57 |
| L8G3W7 | VC83_07867 | *PdBLP3* | 0.071 | 0.001 | -4.15 | 8.03 |
| L8FW73 | VC83_01834 | *PdMRed* | 0.229 | 0.003 | -0.41 | 2.83 |
| L8FZQ4 | VC83_01835 | *PdMFS* | ND | 1.000 | ND | ND |
| L8FWF2 | VC83_01836 | *PdOpDH* | 0.446 | <0.001 | -0.60 | 4.33 |
| L8FXH8 | VC83_01837 | *PdSAM* | 0.323 | <0.001 | 0.67 | 6.18 |
| L8FXD3 | VC83_01838 | *PdP450* | 0.228 | 0.315 | -1.84 | 1.44 |
| L8FT99 | VC83_03096 | MRed | 0.994 | 0.002 | 0.36 | 3.59 |
| L8G150* | VC83_08787 | MRed | 0.69 | 0.025 | -1.17 | 3.16 |

\*Based on Total Intensity

Specifically, under Cu-withholding stress, we find an approximate three-fold reduction in intracellular Cu/Zn-SOD1 (L8FTN0) protein levels and an approximate two-fold increase in cytosolic Mn-SOD3 (L8G5D8) levels. Changes in the extracellular Cu-only SOD4 (L8FYM0) and the mitochondrial Mn-SOD2 (L8G9Z6) proteins are also observed under Cu-restrictive growth conditions, albeit changes in their protein levels are more modest than those of their cytosolic counterparts.

In contrast, Cu-overload appears to affect only Cu/Zn-SOD1 protein levels (Table 6). The approximate two-fold increase (*p* = 0.076) in intracellular Cu/Zn-SOD1 levels may indicate a role in intracellular Cu-buffering^48^. Together, these data suggest that all the SODs enzymes encoded in the *Pd* genome are actively translated and participate in maintaining redox homeostasis under extremes of Cu-stress.

The Cu-responsive gene cluster (CRC) comprises five genes (VC83_01834 - 01838), including two cytosolic and three membrane-associated enzymes ^23^. We detect four of the five predicted CRC proteins in our datasets. Only the protein L8FZQ4, the gene product of VC83_01835, was not detected. The two cytosolic proteins L8FXH8 (VC83_01837; *Pd*SAM) and L8FWF2 (VC83_01836; *Pd*OpDH) are highly regulated by Cu-withholding stress. *Pd*SAM is the third highest ranking DAP based on Log_2_FC (Log_2_FC = 6.18) and displays a more than 70-fold increase in protein levels versus control cells (Table 3); *Pd*SAM also represents the 38^th^ most abundant detectable protein under Cu-restrictive growth conditions (Figure 4c; Table 1).

Whereas *Pd*OpDH is the seventh-highest ranking DAP based on Log_2_FC (Log_2_FC = 4.33) and displays a 20-fold increase in protein levels versus control growth conditions and represents the 961 most abundant detected protein in Cu-starved cells (Table 1, Table 3). The other two CRC proteins, L8FW73 (VC83_01834, *Pd*MRed) and L8FXD3 (VC83_01838, *Pd*P450), are also more abundant in Cu-starved cells. However, their increases are more modest than those of other CRC proteins, with Log_2_FC values of 2.83 and 1.44, respectively. This suggests that CRC enzymes are regulated by Cu-withholding stress and may play an important role in this adaptation response.

#### 3.3.3 Cell surface, secreted and cell wall associated proteins

The fungal cell surface is a vital location at the host-pathogen interface, serving both as a protective barrier against the host and as a compartment for securing essential resources^20^. Under Cu-restriction, fungi rely on high-affinity cell surface CTR transporters to traffic reduced Cu(I) ions across the plasma membrane. CTR trafficking activity can be boosted by the concerted action of cell-surface metalloreductases (MReds) and, in some fungi, homologs to the Cu-scavenging protein BIM1/Cbi1 ^25^ found in *C. neoformans.* The *P. destructans* genome encodes two CTR isoforms, *Pd*Ctr1a (L8FYI1) and *Pd*Ctr1b (L8GBD3). Both CTR1 isoforms are transcriptionally activated by the Cu-withholding stress response^29^. In this study, we find that the *Pd*Ctr1a (L8FYI1) protein shows the most dramatic changes upon Cu-withholding stress, ranking as the second most regulated protein based on Log_2_FC (Log_2_FC = 7.18) and is 145-fold more abundant in BCS treated versus control cells (Table 6). Whereas the *Pd*Ctr1b (L8GBD3) isoform protein also displays high levels of regulation under Cu-withholding stress, with a 59-fold increase (log_2_FC = 5.88) in protein levels in BCS-treated cells compared to control growth conditions (Table 6). This behavior suggests that both isoforms are needed *for P. destructans* to thrive under low-Cu conditions.

The Cu-scavenging protein BIM1/Cbi1 in *C. neoformans*,^25^ has been shown to interact with *C. neoformans* CTR1 to facilitate cellular Cu-import. The *P. destructans* genome encodes three BIM1/Cbi1 proteins, which we have identified as BLP1-3. We were unable to detect unique peptides for BLP1 (VC83_02818; L8FVD4) in our datasets. However, we find that two BLP isoforms, L8FUI0/VC83_03095 (BLP2) and L8G3W7/VC83_07867 (BLP3), are the top two most abundant detected DAPs and they display high levels of enrichment under Cu-withholding conditions (Table 3, Table 1). An 11.9- (Log_2_FC = 3.57) and 260-fold (Log_2_FC = 8.03) increase in BLP2 and BLP3 protein levels, respectively, is detected in BCS-treated versus control cells (Table 1). Notably, *Pd*BLP2 peptide levels are ubiquitous detected in *P. destructans* cultured under all tested growth conditions and are found in the top 4% of all detectable proteins (Supplemental Table S3). This observation may indicate that *Pd*BLP2 may be a core component of the *P. destructans* extracellular matrix. Together, our data are consistent with the notion that the two BLP and the two CTR1 isoforms are abundant in Cu-starved *P. destructans* cells.

We also identify several other secreted or cell-surface-associated proteins that exhibit large and significant alterations in response to Cu-withholding stress (Supplemental Table S6). In this table are several oxidoreductases and carbohydrate-associated proteins DAPs. This includes two flavin adenine dinucleotide (FAD) oxidoreductases (i.e. L8G4Z9, L8FS39), three carbohydrate associated proteins (i.e. L8FS56, L8G482, L8G4K3) and one esterase (L8G8Q9). We interpret that changes in these protein levels may indicate that unique redox-active chemistry is occurring at the cell wall of Cu-starved cells, which may cause distinct fungal cell wall organization or structure.

Metalloreductases (MRed) assist in metal import by supplying electrons to reduce oxidized metal ions (*e.g.,* Cu^2+^ or Fe^3+^) at the cell surface, to facilitate their mobilization across the plasma membrane via selective metal transporters. The *P. destructans* genome encodes 10 predicted MRed genes, of which three (*i.e.,* VC83_01834, VC83_03096, and VC83_08787) have been reported to respond to Cu-stress^23^. We observe only the three Cu-responsive MReds in our global proteomic datasets and, in general, their abundance mimics that observed in transcriptomic studies (Table 6). The gene product of VC83_01834 (L8FW73) is encoded in the CRC and shows a modest increase in protein levels under Cu-withholding stress, with a Log_2_FC of 2.83 relative to control growth conditions. Whereas L8FT99, the protein associated with the gene VC83_03096, displays the largest increase in protein levels upon Cu-withholding stress with a Log_2_FC = 3.59. We also detect peptides corresponding to MRed L8G150, the product of VC83_08787, which shows a Log_2_FC of 3.16 versus control cells. However, due to high sequence similarity between the two MRed proteins L8G150 and L8G2Z0, we cannot definitively assign these changes solely to changes in L8G150 levels. Together, our data suggest that under Cu-restriction, the *P. destructans* cell surface houses at least three MRed proteins that can participate in reductive chemistry at the cell surface. However, the metal ion selectivity and activity of these enzymes warrant future investigation.

### 3.4 P. destructans proteome under chronic Cu-overload stress

In general, chronic Cu-overload results in less perturbations in the *Pd* proteome than Cu-withholding conditions. Homology searches of the 25 most-abundant DAPs upon Cu-withholding stress based on exclusive intensity levels, yield close homologs in *A. fumigatus* but far fewer are detected in the *C. albicans*, *S. cerevisiae*, and *C. neoformans* fungi (Table 2 and Table 4). The top four most detectable DAPs which include L8FSJ0, L8FXZ4, L8G189, and L8G4Z9 all possess predicted secretion signal peptide sequences and encode for cell wall or oxidoreductase proteins. This, in combination with the additional increase in two secreted multicopper oxidase (MCO) enzymes, L8G3T1 and L8G0J2, suggests that extracellular oxidative chemistry might be occurring to alleviate copper overload in the extracellular environment. Other notable changes in intracellular protein levels with identifiable homologs include L8G2W5 (a homolog of a ribosomal protein RPS27A), L8FU21 (a mitochondrial medium chain dehydrogenase/reductase enzyme), L8GB07 (a homolog of the URE2, which is involved in nitrogen and ROS metabolism), L8FX44 (a p450 enzyme involved in sterol metabolism), and L8FWC6 (a subunit of cytochrome *c* oxidase).

We observe that 10 of the 25 DAPs identified under Cu-overload conditions based on total intensity are also represented in 25 DAPs displaying the largest fold changes versus control cells (Table 4). In general, many of the identified DAPs displaying large increases upon copper overload do not have a direct homolog in *S. cerevisiae* or other model fungi. However, L8FW89, a CCC2 homolog, displays a 5.6-fold increase (Log_2_FC = 2.49) in protein levels and may be involved in exporting excess copper from the cytosol into secretory pathway to limit Cu-toxicity. Additionally, a JLP1 homolog (L8G085), is also more abundant in cells experiencing Cu-overload growth conditions with a Log_2_FC = 2.49 vs control conditions. JLP1 is in involved in the sulfur catabolism and increase levels may suggest Cu-overload impacts global sulfur metabolism.

We note that several cytochrome P450 monooxygenase enzymes and DNA associated proteins are identified as significant DAPs under chronic copper overload stress. This includes the four monooxygenases L8G4Y4, L8G0Z6, L8FUQ8, and L8G3R0, which display increased protein levels (Figure 3b, Table 5). However, secondary analysis of these proteins reveals similar increases in protein levels under Cu-withholding conditions, suggesting that these proteins may participate in a general stress response (Supplemental Table S7). We also observe significant reductions in DNA associated proteins L8G8U3 (a homolog to the kinetochore protein NUF2), and L8FPD7 (a SWI5 homolog), L8G614 (a histone methyltransferase homologous to SET1), L8FQ89 (a histone deacetylase homologous to HST2). This may indicate general genomic instability under Cu-overload stress conditions (Figure 3b, Supplemental Table 8). Taken together, our data may indicate Cu-overload stress induces a DNA-damage stress response and may alter general carbon, nitrogen, sulfur and oxygen metabolism.

### 3.5 Validation of anti-sera for detecting Cu-withholding biomarkers

Limited molecular biology tools and knockout strains have been reported in *P. destructans,* and thus the cellular and metal-cell biology of this organism is poorly described ^32, 33^. We sought to use the Cu-stress growth conditions and this proteomics dataset to validate metal-responsive genes that could serve as biomarkers for *P. destructans* Cu-withholding stress. Although *Pd*BLP2 and *Pd*BLP3 are highly abundant in Cu-starved cells, they may not be ideal protein targets because they are predicted to be GPI-anchored proteins,^23, 29^ which may exhibit complex extracellular trafficking or reorganization patterns at the cell surface^49^. Thus, we turned to identify two novel intracellular and membrane-associated proteins. We identified the candidate proteins L8FYI1 (*Pd*Ctr1b isoform) and three members of the CRC [L8FW73 (*Pd*MRed); L8FWF2 (*Pd*OpDH); L8FXH8 (*Pd*SAM)]^23^ which are regulated DAPs which respond to Cu-withholding stress. *Pd*Ctr1b and *Pd*MRed are membrane proteins, whereas *Pd*OpDH and *Pd*SAM are predicted to reside in the cytoplasm. Together, these proteins would be ideal candidates for probing biochemical processes in whole-cell lysates. Figure 5 displays whole-cell lysate western blots against these candidate protein targets, as well as the previously described *Pd*CTR1a protein^29^. We observe a strong correlation between the quantitative peptide quantification abundance profiles and western immunodetection against *Pd*Ctr1a, *Pd*Ctr1b, *Pd*OpDH, and *Pd*SAM protein targets (Figure 5). However, broad, poorly resolved high-molecular-weight peaks are observed for *Pd*MRed (Figure 5c). This behavior may indicate high glycosylation at one or more of the four predicted *Pd*MRed asparagine residues (i.e., N52, N153, N295, N325), making it less suitable as a biomarker for Cu-withholding stress. Our observations, using immunoblot detection of *Pd*CTR1a, *Pd*CTR1b, *Pd*SAM, and *Pd*OpDH, in combination with our global proteomics, suggest that these protein targets may represent promising new biomarkers in *Pd* samples for chronic Cu-withholding stress for western blot or proteomic analyses.

**Figure 5.**
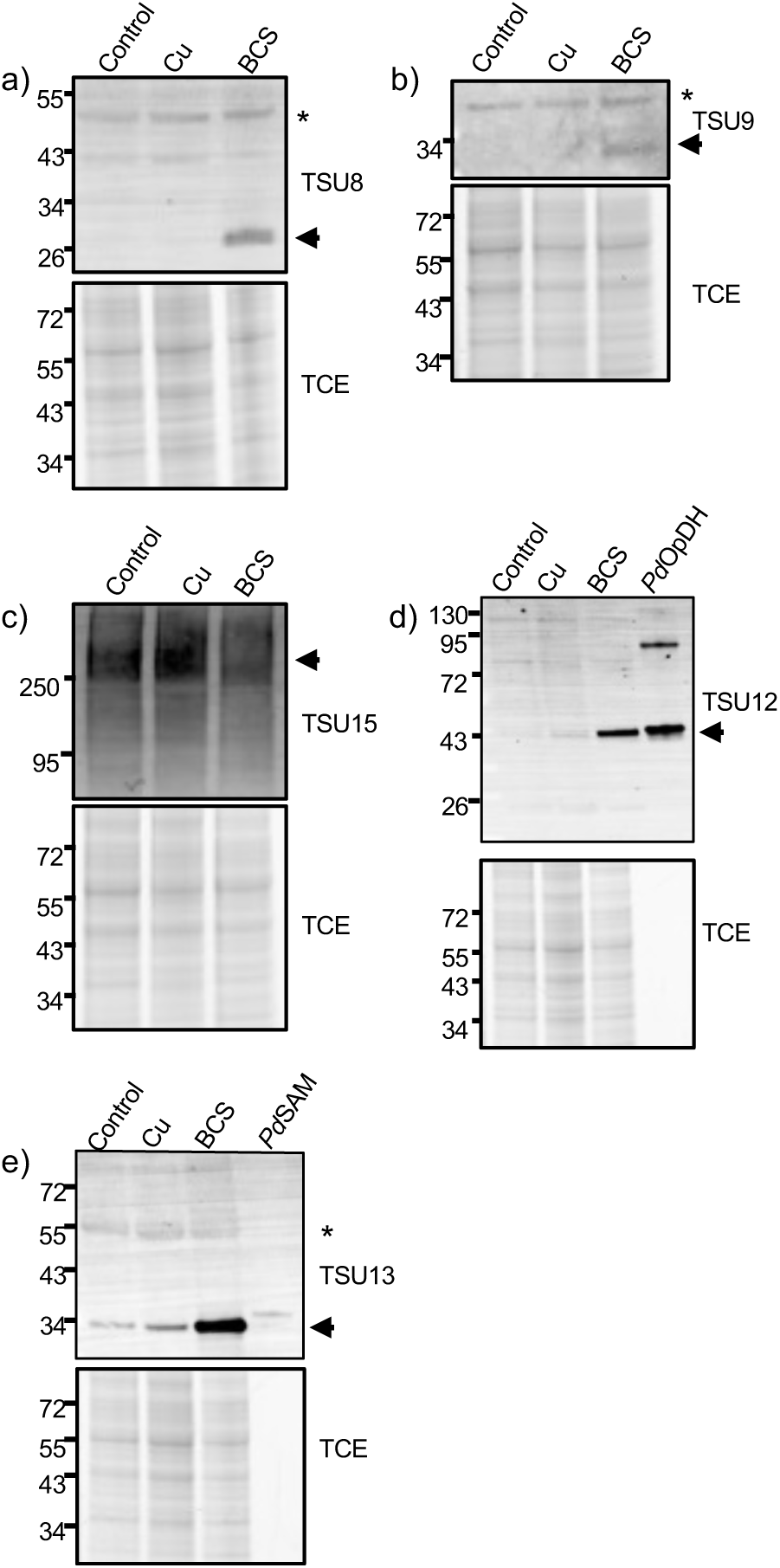
Western immunoblots of *P. destructans* cell lysates against *Pd*CTR1 isoforms and *Pd*CRC proteins. Western blots for (a) *Pd*CTR1a (L8FYI1; VC83_00191) (b) *Pd*CTR1b (L8GBD3; VC83_04814) (c) *Pd*MRed (L8FW73; VC83_01834), (d) *Pd*OpDH (L8FWF2; VC83_01836), (e) *Pd*SAM (L8FXH8; VC83_01837) are displayed in the top panel. Below each western blot is a TCE-stained loading control of the ∼25 – 90 kDa region of the SDS-PAGE gel. Purified *Pd*OpDH (10 ng) and *Pd*SAM (10 ng) are used as loading standards in panels (d) and (e), respectively. Non-specific bands are indicted with * symbol.

## 4. Discussion

*Pseudogymnoascus destructans* and the infectious disease WNS in bats are important to both environmental and human public health^50^. Currently, there is a fundamental gap in our understanding of the mechanisms by which this pathogen adapts and thrives in its infected bat host and across diverse environmental niches. Previous RNA-seq studies at active WNS-infection sites on bats suggest that *P. destructans* is starved of trace-metal micronutrients ^21, 22^. Complementary laboratory growth studies under copper-restrictive conditions have revealed that several putative virulence factors associated with the WNS-disease state can be induced under Cu-withholding stress^23^. Thus far, methods to understand *P. destructans* basic metabolic and cell biology have been primarily focused on growth and transcriptional studies,^21–23, 51^ with gaps in understanding *P. destructans* cell biology at the protein level. This report focuses on the use of quantitative global proteomics in *P. destructans* to identify adaptations to Cu-stress environments under laboratory culture conditions.

We find that approximately 48% of the predicted *P. destructans* proteome is detected in laboratory-cultured samples. This breadth of expressed proteins is comparable to that observed for global proteomic studies in *C. neoformans*^52^ but approximately two-fold higher than for *C. albicans*^53^ or *S. cerevisiae*^54^. This suggests that much of the approximately 9000 predicted *P. destructans* proteome is transcriptionally accessible or actively translated during laboratory culture conditions.

How does *P. destructans* adapt to Cu-restrictive stress? This adaptation may be necessary during cultivation in cave environments to outcompete other microbes living on scarce resources or to circumvent host nutritional immunity defense mechanisms that restrict metal bioavailability^6, 55^. Indeed, previous transcriptional studies at active WNS disease sites have shown elevated levels of potent host metal-binding proteins, including members of the S100 family^21^. Under laboratory culture conditions, it is apparent that *P. destructans* is well-suited to adapt to and efficiently propagate in Cu-restrictive environments by using two active extracellular metal-scavenging pathways and prioritizing intracellular Cu-metalloenzyme levels. First, Cu limitation leads to large increases in cell-surface Cu transporters and extracellular Cu-scavenging proteins. Both Cu-transporters *Pd*Ctr1a and *Pd*Ctr1b are detected in high abundance and at comparable levels. At the same time, protein levels of the two BLP isoforms, *Pd*BLP2 and *Pd*BLP3, are detected at higher levels. However, the relative levels of these BLP isoforms across different Cu-stress conditions appear distinct. *Pd*BLP2 levels remain high across all copper stress conditions, whereas *Pd*BLP3 protein levels appear to be more restricted to cells experiencing Cu-restrictive growth conditions. This suggests that *Pd*BLP2 and *Pd*BLP3 may play unique roles in Cu-fungal biology. *Pd*BLP2 and *Pd*BLP3 may interact with a specific *Pd*CTR isoform or regulate CTR permease activity. Under chronic Cu-limiting stress, *C. neoformans* expresses two CTR transporters (*CTR1* and *CTR4*) variants^56^. However, *C. neoformans* BIM1/Cbi1 only participates in relaying copper to the CTR1 isoform. More recent studies with *C. neoformans* BIM1/Cbi1 suggest that BIM1/Cbi1 may also participate in maintaining fungal cell wall chitin/chitosan integrity.^26^ With two highly abundant BLP isoforms present at the *P. destructans* cell surface, specialized roles in Cu-scavenging and maintaining fungal cell wall integrity are possible. Many fungal genomes encode BLP isoforms, and future studies should be directed at determining how individual BLP isoforms contribute to fungal fitness.

The second potential copper-scavenging pathway may involve a small-molecule metabolite produced and secreted by proteins encoded in a previously identified gene cluster, the *Pd* Cu-responsive gene Cluster (CRC).^23^ This study provides evidence both through quantitative proteomics and immunoblot assays that *Pd*SAM and *Pd*OpDH protein expression levels are good indicators of *Pd* cells experiencing chronic Cu-withholding stress. While the biochemistry and fungal cell biology of the *Pd*CRC are not yet well defined, its behavior at both the transcriptional and translational levels strongly implicates a role in metal ion homeostasis^23^. As biochemical and molecular biology tools for *P. destructans* advance, there will be opportunities to further interrogate extracellular Cu-scavenging pathways involving *Pd*BLP isoforms and the *Pd*CRC.

*Pseudogymnoascus destructans* may avoid using abundant, intracellular Cu-utilizing enzymes under Cu-restrictive conditions to maintain efficient propagation. This is supported by changes in the levels of the most abundant copper enzymes in the mitochondria and cytoplasm, CcO and Cu/Zn SOD, respectively. Upon Cu-restrictive growth, many significant alterations in the levels of CcO subunits and chaperones are observed, accompanied by an increase in alternative oxidase (AOX) protein levels. Similar modifications in CcO and AOX activities have been observed in *C. albicans* under Cu-limited conditions, enabling efficient respiratory growth under Cu restriction^57^. Additionally, copper restriction leads to a decrease in intracellular Cu/Zn-SOD levels and an increase in the two Mn-containing SODs, presumably in the cytoplasm and mitochondria. This reciprocal regulation in the distribution of Cu- and Mn-SOD levels is observed in *C. albicans* under stationary growth conditions^58^ as well as under Cu-limitation and infection conditions^59^. This switch in intracellular SOD levels may be favored to preserve cellular SOD activity and divert trace Cu ions to CcO or Cu-MCOs to facilitate Fe-import under Cu-scarcity^60–62^. In *C. albicans* and *C. neoformans,* this cytosolic Mn-SOD regulation is controlled at the transcriptional level by a copper-sensing transcription factor^59, 63^. A similar mechanism involving an unidentified Cu-sensing transcription factor may also occur in *P. destructans,* as the transcriptional changes in Cu/Zn-SOD and cytosolic-Mn SOD levels grown under similar Cu-stress growth conditions also correlate with this study^23^. Based on our observations of culturing *P. destructans* under varying copper stress conditions, we proposed a model of significant changes in the proteomes of extracellular, intracellular, and mitochondrial proteins under Cu-withdrawal and Cu-overload conditions (Figure 6).

**Figure 6.**
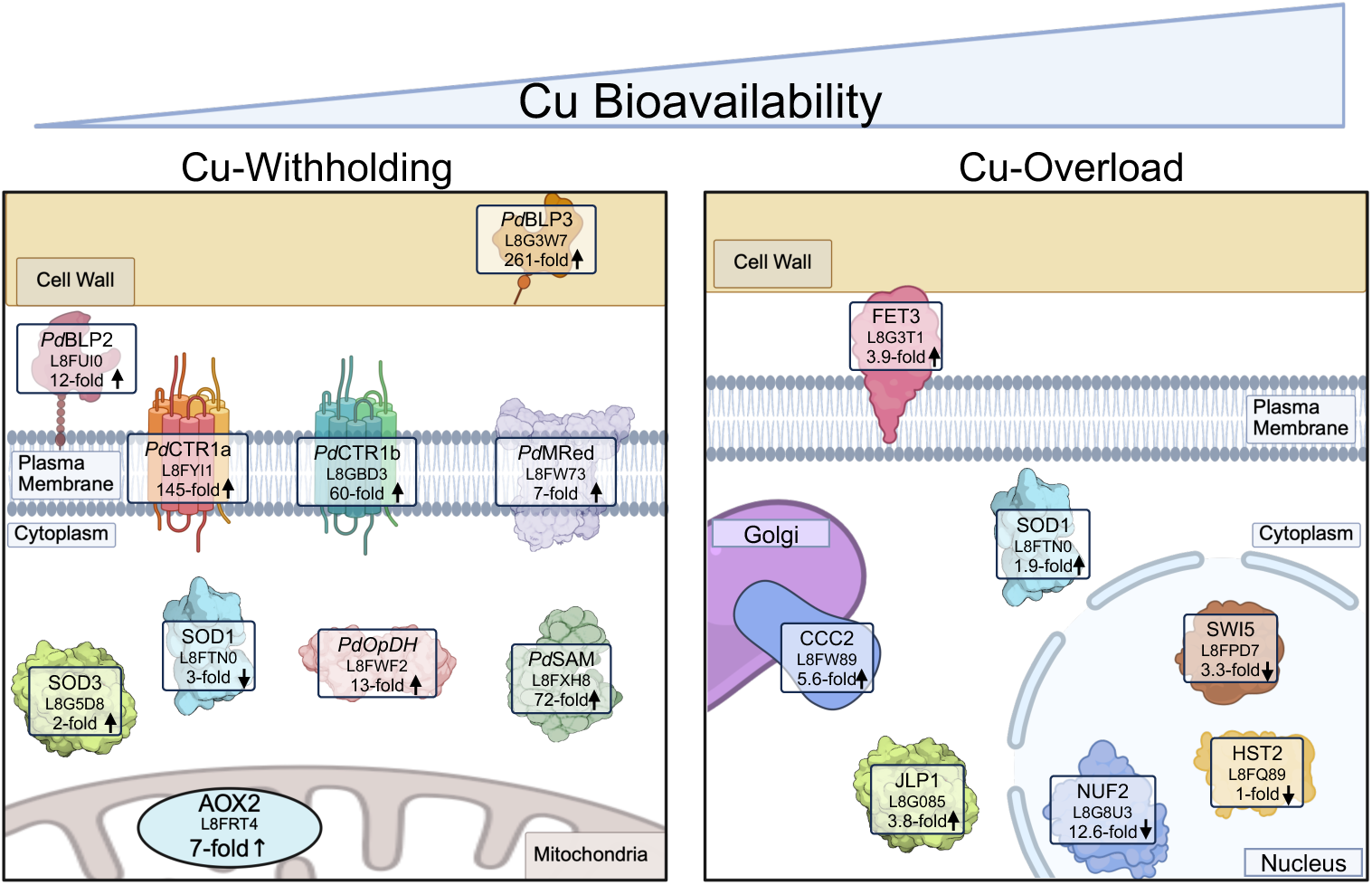
Model of select *P. destructans* proteins that are significantly altered under Cu-withholding and Cu-overload stress.

Previous transcriptomic studies in *P. destructans* have indicated that high Cu-exposure can lead to increased transcript levels of genes associated with DNA damage and replication stress^23^. Our data in this report are broadly consistent with these observations at the protein level. We note that the *P. destructans* genome encodes for a histone H3 variant (L8FY77) possessing an analogous H3-C110;H113 Cu-binding site and is predicated to have intracellular copper reductase activity^64^. This feature is conserved in higher eukaryotes but is absent in many fungal species^65^. How this histone feature may impact *P. destructans* genome stability and cellular redox homeostasis, especially under high copper stress conditions, is unknown but warrants further study.

There are limited molecular biology tools established for modifying the *P. destructans* genome^33^ and validated reagents for studying *P. destructans* cell biology. Previously, studies have reported antibodies against the secreted subtilisin-like serine protease *Pd*SP1^66^ and the high-affinity Cu-transporter *Pd*Ctr1a^29^. Here, we also describe the development and validation of anti-sera for *P. destructans*-specific proteins that respond to Cu-withholding stress. The addition of antisera against Cu-responsive protein targets, including cytosolic *Pd*SAM and *Pd*OpDH, as well as the cell surface *Pd*Ctr1b Cu-transporter, will provide opportunities to further interrogate *P. destructans* cell biology in both *in vitro* and infection settings.

While we have demonstrated that a substantial portion of the *P. destructans* predicted global proteome can be detected, this study has some limitations. First, our design used a single temperature, growth media, and time point. Previous studies have shown that *P. destructans* transcriptional profiles are sensitive to growth media composition and temperature^67, 68^, and thus the proteome may also be sensitive to such effects. This study was performed at 15 °C near the optimal *P. destructans* growth temperature of 15.8 °C on chemically defined media to facilitate high levels of fungal replication^69^. However, the temperature of hibernating bats can vary with long periods of low, near ambient temperature, with short arousal intervals with skin temperatures near 30 °C ^70^. Second, this current study focused on quantifying total protein levels, which may not accurately reflect properly metalated proteins or levels of metalloenzyme activity. The total cellular activity levels of Cu/Zn SOD and CcO may be substantially lower than quantified by our mass spectrometry-based methods. Further studies into the relative activities and quantities of CcO, AOX, Mn-SOD, and Cu/Zn-SOD will be insightful for validating relative Cu-loading levels. Third, in this study, we were unable to detect peptide fragments for metallothionein or the COX17 CcO Cu-chaperone, which are known to participate directly in Cu-binding and trafficking under Cu-stress conditions in fungi^71–73^. We anticipate that the inability to detect peptides from these critical players in Cu-biology could be a limitation of our proteomics workflow^74^. Finally, many of the DAPs identified under copper-overload conditions have minimal direct homologs in well-characterized model fungi. We anticipate that, as more proteomic and transcriptomic studies are conducted in other *Pseudogymnoascus* species or related filamentous fungi, we will gain insight into the metal stress response of this unique organism.

This is particularly true for *P. destructans* exposed to high Cu-levels.

## 5. Conclusions

Our study provides a snapshot of the *P. destructans* proteome and its adaptation to copper micronutrient withholding and overload stress. We find that Cu-withholding stress leads to substantial modification of the *P. destructans* proteome, in that there is reprioritization of high-utilizing Cu-proteins and Cu-acquisition pathways. Specifically, Cu-containing SOD levels, as well as the respiratory enzymes cytochrome *c* oxidase and alternative oxidase pathways, are highly impacted. Future studies warrant investigation of the activities of these enzymes under variable Cu-stress and temperature conditions. On the other hand, chronic Cu-overload exposure is readily tolerated by *P. destructans,* resulting in minimal impacts on the global protein levels. Because high Cu-exposure causes alterations in select DNA associated proteins and basic carbon-, nitrogen- and sulfur-metabolic proteins, further investigation into these pathways is warranted. Together, this *P. destructans* global proteomic dataset and the antibodies developed in this study to monitor the *P. destructans* metal stress response will serve as valuable resources to advance understanding of *P. destructans* cellular and metal-cell biology.

Raw and processed data files have been deposited in the MassIVE repository (massive.ucsd.edu) under accession number [MSV000101104] and in ProteomeXchange repository under accession number PXD[PXD075515].

## Author Contributions

Conceptualization, S.T.W. and R.L.P; Methodology, A.D.F., S.A., S.T.W., Y.L. and R.L.P; Validation, A.D.F., S.T.W., Y.L., and R.L.P; Formal analysis, A.D.F., S.A., S.T.W., Y.L., and R.L.P; Investigation, A.D.F., S.A. and R.L.P; Resources R.L.P.; Data curation, A.D.F., S.T.W., and R.L.P; Writing—original draft, A.D.F. and R.L.P; Writing—review & editing, A.D.F., S.A., S.T.W., Y.L. and R.L.P; Visualization, A.D.F., S.T.W., Y.L., and R.L.P. All authors have read and agreed to the published version of the manuscript.

## Data Availability Statement

The data presented in this study are available from the corresponding author upon reasonable request.

## Acknowledgments

A.D.F., S.A. and R.L.P. would like to thank the Texas State University for financial support. Mass spectrometry analyses were conducted at the University of Texas Health Science Center at San Antonio (UTHSCSA) Institutional Mass Spectrometry Laboratory, with expert technical assistance of Sammy Pardo and Dana Molleur, under the direction of S.T.W., supported in part by UTHSCSA and by the University of Texas System Proteomics Core Network for purchase of the Orbitrap Fusion Lumos mass spectrometer.

---

**Supplemental File 1.** Supplemental Figure S1. Microscopy images of *P. destructans* spores isolated from Control, Cu-Overload, and Cu-Withholding growth conditions.

**Supplemental File 2.** Supplemental Table S1: Table of serum products used for the detection of *Pd* DAPs responding to Cu-withholding stress.

**Supplemental File 3.**

Supplemental Table S2. DIA-MS processed data

Supplemental Table S3. Significant DAPs identified under Cu-withholding stress

Supplemental Table S4. Significant DAPs identified under Cu-overload stress

Supplemental Table S5. DAPs used to generate the Venn diagram

Supplemental Table S6. DAPS with predicted secretion signals displaying large changes under Cu-withholding stress

Supplemental Table S7. Table of annotated P450 proteins under Cu-stress conditions

Supplemental Table S8. DAPs displaying the highest FC reduction under Cu-overload stress

**Supplemental Table S1.** Table of serum products used for the detection of *Pd* DAPs responding to Cu-withholding stress.

| Serum Identifier | *Pd* Protein Target | Peptides/Protein | Western Blot Dilution Ratio |
| --- | --- | --- | --- |
| TSU8<sup>1</sup> | VC83_00191 | KARQEARWLDCEMHRRY<u>C</u>;<br>GKPGLRERVALHKDAK<u>C</u> | 1:500 |
| TSU9 | VC83_04814 | MSHSMGHGDHDASAAR<u>C</u>;<br>YDAALLKRRDELPHEELA<u>C</u> | 1:1000 |
| TSU12 | VC83_01836 | Recombinant full-length protein,<br>VC83_01836, Plasmid pAS33 | 1:1000 |
| TSU13/14 | VC83_01837 | Recombinant full-length protein,<br>VC83_01837, Plasmid pAS32 | 1:1000 |
| TSU15 | VC83_01834 | <u>C</u>NFLLPKDEV DGIEYS;<br><u>C</u>PIARSDSATREEEAK | 1:1000 |
| TSU17/18 | VC83_01835 | <u>C</u>GRSEKGQVEGGSNSY;<br><u>C</u>LVKRAASGKIITPED | 1:1000 |

For peptide fragments, the reactive cystine used for KLH conjugation is underlined. <sup>1</sup> As previously reported by Anne, S.; Friudenberg, A.D.; Peterson, R.L. Characterization of a High-Affinity Copper Transporter *CTR1a* in the White-Nose Syndrome Causing Fungal Pathogen *Pseudogymnoascus destructans*. *J. Fungi* **2024**, *10*, 729.

**Supplemental Figure S1.**
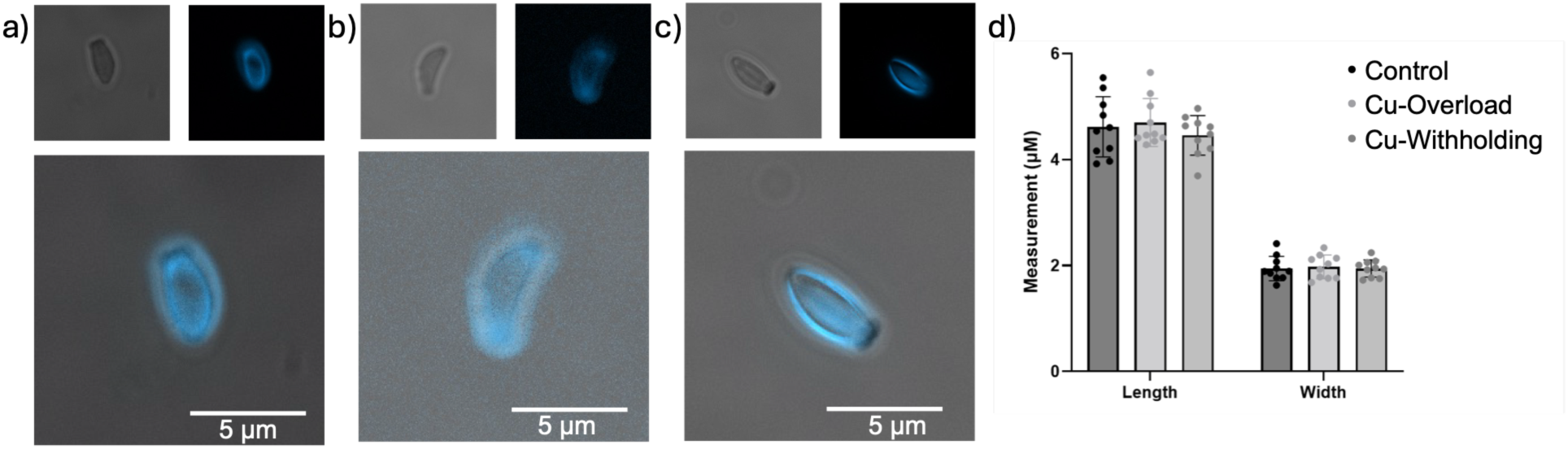
Microscopy images of *P. destructans* spores isolated from Control, Cu-Overload, and Cu-Withholding growth conditions. A-C images of *P. destructans* Cells imaged under bright field (top left), Calcofluor-white (top right), and merged (bottom). Samples were cultured under (a) Control, (b) Cu-Overload, (c) Cu-Withholding growth conditions. (d). Graph displaying spore morphology parameters under different growth conditions, n = 10.

