## Supplemental File 1 for "The *Pseudogymnoascus destructans* Proteome Under Copper Stress Conditions"

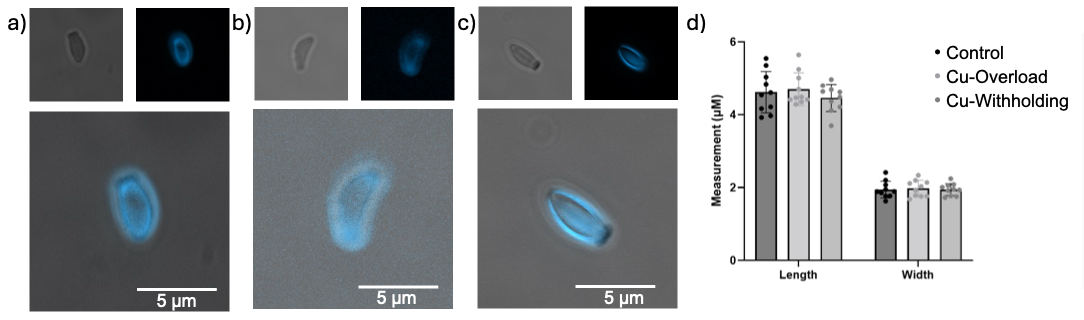

**Supplemental Figure S1**. Microscopy images of *P. destructans* spores isolated from Control, Cu-Overload, and Cu-Withholding growth conditions. A-C images of *P. destructans* Cells imaged under bright field (top left), Calcofluor-white (top right), and merged (bottom). Samples were cultured under (a) Control, (b) Cu-Overload, (c) Cu-Withholding growth conditions. (d). Graph displaying spore morphology parameters under different growth conditions, n = 10.
