## Supplemental File 2 for "The *Pseudogymnoascus destructans* Proteome Under Copper Stress Conditions"

Supplemental Table S1. Table of serum products used for the detection of *Pd* DAPs responding to Cu-withholding stress.

| **Serum Identifier** | ***Pd P*rotein Target** | **Peptides/Protein** | **Western Blot Dilution Ratio** |
| --- | --- | --- | --- |
| TSU8^1^ | VC83_00191 | KARQEARWLDCEMHRRYC; GKPGLRERVALHKDAKC | 1:500 |
| TSU9 | VC83_04814 | MSHSMGHGDHDASAARC; YDAALLKRRDELPHEELAC | 1:1000 |
| TSU12 | VC83_01836 | Recombinant full-length protein, VC83_01836, Plasmid pAS33 | 1:1000 |
| TSU13/14 | VC83_01837 | Recombinant full-length protein, VC83_01837, Plasmid pAS32 | 1:1000 |
| TSU15 | VC83_01834 | CNFLLPKDEVDGIEYS;  CPIARSDSATREEEAK | 1:1000 |
| TSU17/18 | VC83_01835 | CGRSEKGQVEGGSNSY; CLVKRAASGKIITPED | 1:1000 |

For peptide fragments, the reactive cystine used for KLH conjugation is underlined. ^1^ As previously reported by Anne, S.; Friudenberg, A.D.; Peterson, R.L. Characterization of a High-Affinity Copper Transporter *CTR1a* in the White-Nose Syndrome Causing Fungal Pathogen *Pseudogymnoascus destructans*. *J. Fungi* **2024**, *10*, 729.
